# Climate change has desynchronized insect and vegetation phenologies across Europe

**DOI:** 10.1101/2023.12.11.571152

**Authors:** Yanru Huang, Chaoyang Wu, Wenjiang Huang, Yang Liu, Man Qi, Jie Bai, Yingying Dong, Samuel J L Gascoigne, Philippe Ciais, Josep Peñuelas, Roberto Salguero-Gómez

## Abstract

Climate change is drastically affecting the phenologies of species worldwide, including shifts in development^1–3^. The potential impact of climate change on the synchronicity of plant-insect phenology is particularly concerning since the stability of pollination networks and food chains depends on their tight temporal dependencies^4–6^. Furthermore, the recently reported “insect armageddon”^7^ makes it urgent to establish trends and identify primary drivers of plant-insect phenological synchrony. Here, coupling high-resolution remote sensing and citizen science data across Europe over 34 years, we examine the differences in occurrence dates of 1,584 herbivorous insects across four orders and the corresponding dates of leaf unfolding. We find that 61.2% of the vegetation and insect phenologies have become highly asynchronous, with vegetation phenology advancing four-fold faster than insect phenology. These trends were modulated by insects’ life-cycles and taxonomic order. A primary driver of this phenological mismatch is the higher sensitivity of vegetation phenology than insect phenology to climate, which has prevented insects from matching the pace of plant phenological advance in the growing season. Our analyses provide a unique continental overview and underlying mechanisms of the asynchronicity between vegetation and insect phenologies, thus enhancing our ability to predict and manage its potential cascading ecological effects.

## Main

The timing of events during the life cycle of an organism (e.g., maturation, reproduction, and dispersal) is fundamental to its persistence^8^. Species phenologies are key to the maintenance of entire communities^9^ and their ecosystem services^10^. Important phenological shifts, however, have been reported worldwide in recent decades^11^. For example, global warming has advanced the date of laying of great tits (*Parus major*) by more than two weeks in the last six decades^12^, and the start of the growing season on the Qinghai-Tibetan Plateau has advanced by nearly 10 days in the last two decades^13^.

Phenological shifts due to climate change are thought to optimise species fitness, or at least to reduce the negative impacts of a changing environment^14^. Not all phenological shifts, however, are equal, nor do they have the same consequences. The direction and intensity of phenological shifts vary considerably across taxa^15^, locations^16^, and environments^17^. Importantly, species within a community can have different phenological shifts in response to the same climatic driver, thereby disrupting food webs^18^. Climate change consequently has the potential to impede the temporal and spatial synchrony of biological interactions, ultimately leading to trophic collapse^19^.

The phenological synchrony between vegetation and insects at a given location ensures the match between the supply and demand of resources. This temporal match is crucial to maintain the flow of energy in food chains, which ultimately support viable populations and biodiversity^20^. Insect feeding is tightly linked to specific stages of vegetation development because this relationship ensures that insects obtain the required resources^21^. Phenological asynchrony emerges when the phenologies of co-existing vegetation and insects shift at different rates and/or in different directions^22^. The ecological consequences of this phenomenon can be devastating, including mismatches in food-pollinator interactions^23^ or the decline in insect biodiversity^24,25^. Indeed, 17-50% of pollinators are projected to experience interruptions in food supply due to phenological mismatches under the on-going gradual warming^26^. Eighty-four percent of crops cultivated in Europe directly depend on insect-mediated pollination for their yield^27^, so a potential mismatch between the phenologies of vegetation and insects poses an important threat to the stability of both ecosystems and livelihoods.

The impacts of climate change on the synchrony of vegetation and insect phenologies have been debated for decades. Some studies suggest that climate change has negative impacts on the synchrony of vegetation and insect phenologies^28,29^, but others propose that some species may adapt to climate change and maintain the temporal synchrony of their biological events^21,30^. Approaches to examine this mismatch have predominantly used manipulative experiments and ground-based observations^31,32^. However, we still lack a generalized view about how climate change affects phenological synchrony across large scales and diverse taxa, because findings remain specific to species, geography, and experiment^33^. Remote-sensing studies have verified the widespread advance of vegetation phenology in spring and its delay in autumn in recent decades^34^. The main controversy is about insect phenology, which is characterized by greater complexity and dynamism than vegetation phenology. This complexity challenges our efforts to monitor insect phenology. Fortunately, data from citizen scientists provide a valuable resource for the extensive, long-term, and multispecies insect monitoring^35^. Integrating data from multiple sources can finally allow a comprehensive assessment of the synchrony between vegetation and insect phenologies.

Here, we investigate the trends and driving factors of phenological (a)synchronies between vegetation and insects across Europe from 1982 to 2015, supported by high-resolution remotely sensed data and citizen science. Specifically, (1) we quantify the rate and direction of shifts in vegetation phenology, insect phenology, and their differences (vegetation-insect phenological difference, VID) at both continental and regional scales across Europe. We test four hypotheses about the trends of changes in VID (Figure 1). (2) We next evaluate whether the rate and direction of changes in VID differs amongst insect orders (Hemiptera, Hymenoptera, Coleoptera, and Lepidoptera) and life-cycle voltinism (i.e., number of generations produced per year), and how these differences manifest themselves across latitudinal gradients. Finally, (3) we analyze the differences in the sensitivity of vegetation and insect phenologies to changes in environmental factors. We identify the likely environmental factors driving the asynchrony in VID by combining these analyses with the trends of climate change.

**Figure 1.**
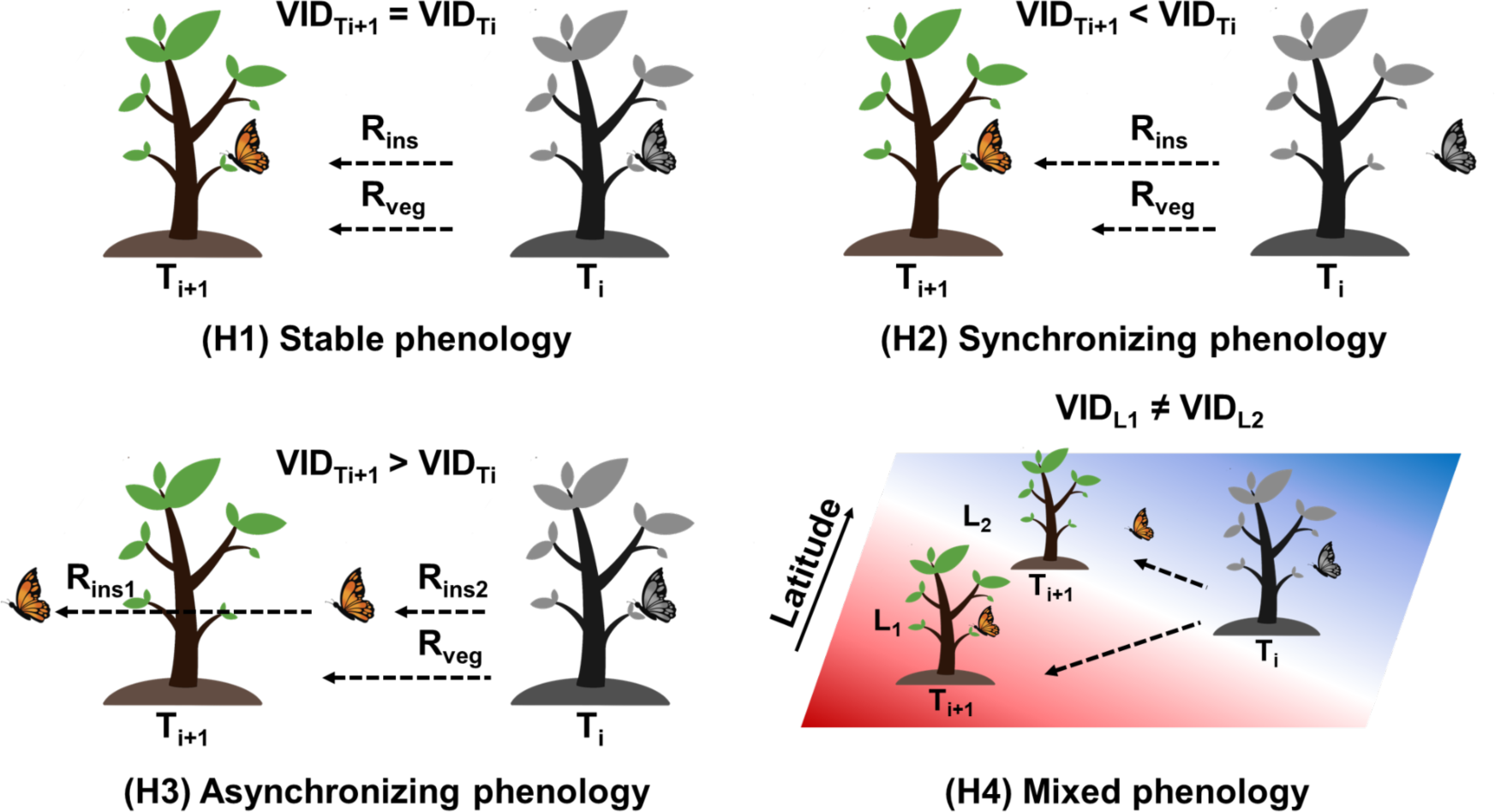
We test four hypotheses regarding how climate change may have shaped vegetation-insect phenological differences (VID) across Europe in recent decades. **H1** - **stable phenology**: the rate of advance (R_veg_ and R_ins_) does not differ significantly between vegetation phenology (leaf-unfolding date, LUD) and insect phenology (insect occurrence date, IOD), so the change between the initial state VID_Ti_ (grey tree and grey butterfly) and VID_Ti_ (coloured tree and butterfly) would not be large. **H2** - **synchronising phenology**: the peak of the phenology of insects starts after the peak of the phenology of vegetation, but both synchronise so that VID_Ti+1_ < VID_Ti_, because R_veg_ < R_ins_. **H3** - **asynchronising phenology**: VID_Ti+1_ is larger than VID_Ti_, with two potential scenarios: ① R_veg_ > R_ins1_, and ② R_veg_ < R_ins2_. **H4** - **mixed phenology**: the spatial trends of VID differs latitudinally due to the spatial heterogeneity of climate change. For example, location L_1_ and location L_2_ would initially have the same VID at time T_i_, but VID_L1_ and VID_L2_ would differ significantly at time T_i+1_. Both LUD and IOD advance as climate change progresses. The background colour in panel H4 represents the spatial heterogeneity of the climatic factors.

### Phenological asynchrony between vegetation and insects has intensified

To establish trends and identify putative mechanisms in the phenological (a)synchronisation between vegetation and insects in Europe, we acquired 34 years of insect occurrence date (IOD) data for 1,584 species of herbivorous insects, observed by citizen scientists, and extracted the leaf-unfolding date of vegetation (LUD) at the corresponding spatiotemporal locations using remote sensing data (Figure S1). Using a quantile-regression model, we found that vegetation phenology LUD in Europe advanced by 12 days during 1982-2015 (-0.37 d/y ± 0.006 S.E.; Table S1), consistent with the estimate of vegetation phenology in spring in northern Europe^36^. In contrast, while insect phenology IOD also significantly advanced during the same period, it did so at a pace four-fold slower (-0.09 d/y ± 0.007) than the vegetation (Wilcoxon V = 199622, *P*<0.001) (Figure 2c). This lag in the advance of insect phenology relative to vegetation suggests a potential decoupling in trophic interactions^37^. Of the 1,807 insect phenological patterns, 79.6% had an increase in VID (with 61.2% being significant, *P*<0.05), only 20.4% exhibited a decrease in VID (with 7.4% significant, *P*<0.05), indicating an exacerbation of phenological asynchrony (H3 scenario ① in Figure 1) at a rate of nearly three days per decade (0.28 d/y ± 0.009; Figure 2d).

**Figure 2.**
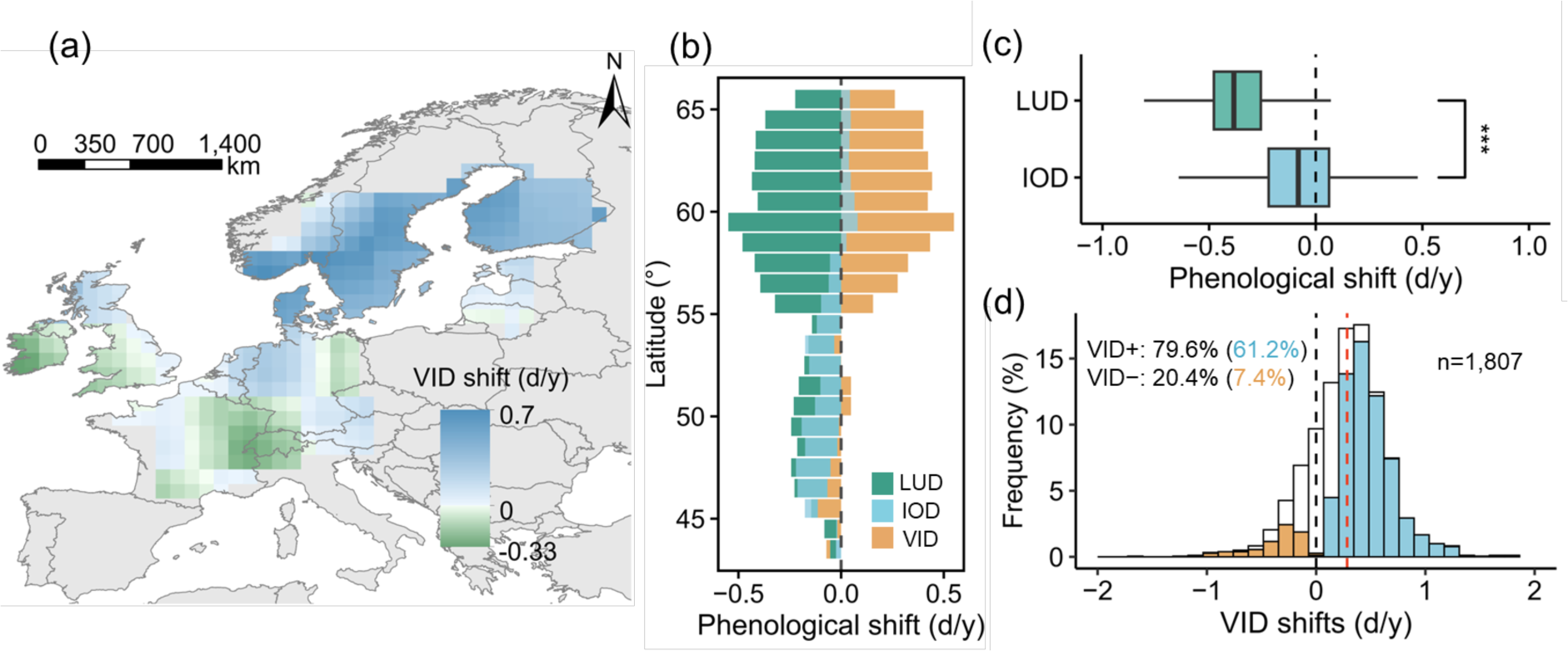
Synchronisation changes between the European leaf-unfolding date (LUD) of vegetation and the insect occurrence date (IOD) (1982-2015). (a) Spatial distribution of the rate of the shift in the vegetation-insect phenological difference (VID) at the regional scale using the moving window algorithm, with green and blue corresponding to synchronous and asynchronous matches in phenology, respectively. (b) Regional trends of LUD (green), IOD (blue), and VID (orange) with phenological shifts as a function of latitude. IOD shifts from advanced to slightly delayed, and LUD trends to increasingly advance as latitude increases. (c) Continental trends of phenological shifts in LUD and IOD, indicating how the phenology of vegetation advances more rapidly than the phenology of insects; ***, *P*<0.001, by a Wilcoxon signed-rank test. (d) The continental trend indicates that the VID of more species patterns is becoming asynchronous, where blue and orange indicate significant asynchronicity (VID+) and synchronicity (VID−), respectively, with white indicating no significant change. The red dashed vertical line represents the mean rate of shift in VID.

The regional rate of change in VID differed significantly between low latitudes (L_1_: ≤55°N) (-0.02 d/y ± 0.004) and high latitudes (L_2_: >55°N) (0.38 d/y ± 0.007) (Wilcoxon W = 1191, *P*<0.001) (Figure 2a, b, Table S2). The expected increase in VID was 0.03 d/y ± 0.001 for each degree increase in latitude. This pattern was attributed to the increased difference in the rates (R) of change of LUD and IOD between the high latitudes (R_LUD_ = -0.42 d/y ± 0.005, R_IOD_ = 0.02 d/y ± 0.003) and low latitudes (R_LUD_ = -0.19 d/y ± 0.005, R_IOD_ = -0.16 d/y ± 0.004). Our analysis also found that VID intensified phenological asynchrony along the latitudinal gradient, consistent with previous observations^5,17^, as illustrated in panel H4 in Figure 1.

### Rate of shift in the vegetation-insects phenological difference varies with taxonomic order and degree of voltinism

The shifts in VID across Europe in the last three decades indicated a strong specificity of insect order (Kruskal-Wallis χ^2^(3) =158.26, *P*<0.001) (Figure 3a, Table S3). For example, the rate of shift of VID during the study period was fastest for Lepidoptera (e.g., butterflies and moths), at 0.34 d/y ± 0.01. This order was followed by Hemiptera (true bugs), with a shift in VID of 0.16 d/y ± 0.05, and by Hymenoptera (e.g., bees), with a shift of 0.15 d/y ± 0.04. In contrast, the rate of shift was slower for Coleoptera (beetles), at only 0.04 d/y ± 0.03. These findings suggested that Coleoptera adjusted their IOD (R_IOD_ = -0.27 d/y ± 0.02) most rapidly to match the changes in vegetation phenology, leading to fewer phenological patterns with significant VID shifts (27.72% *P*<0.05). Conversely, the phenology of Lepidoptera, with a slower rate of adaptation (R_IOD_ = -0.04 d/y ± 0.01), increasingly differed from vegetation phenology (78.23% *P*<0.05), consistent with recent findings for the mean changes in flight day amongst insect orders in Europe^6^.

**Figure 3.**
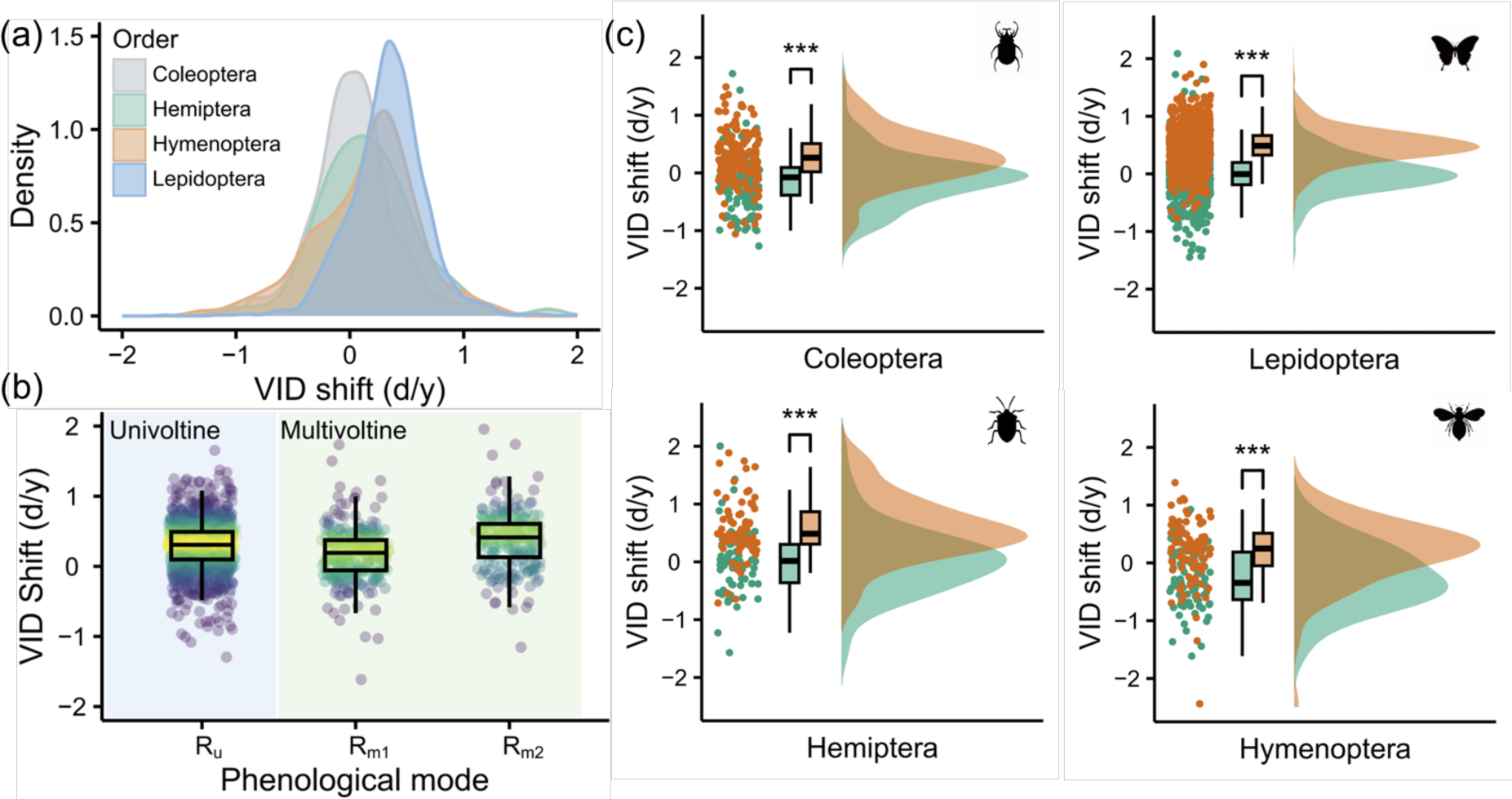
Shift in vegetation-insect phenological difference (VID) across insect orders and levels of life-cycle voltinism (i.e., number of generations/year), with beetles and multivoltine first-generation insects being more capable of tracking shifts in vegetation phenology. (a) Probability density curves of VID shifts across four insect orders. (b) VID shifts in univoltine insects (R_u_) and multivoltine insects (first generation (R_m1_), second generation (R_m2_)). The colours represent the density of data, with yellow and blue indicating high and low densities, respectively. (c) Differences in VID shifts along a latitudinal gradient in Europe for the four insect orders, distinguishing between low (green: L_1_≤ 55°N) and high (orange: L_2_ > 55°N) latitudes. ***, *P*<0.001 by a Wilcoxon signed-rank test.

Further analyses identified significant differences in the rate of variation of VID for each insect order along the latitudinal gradient (L_1_ low and L_2_ high) (Figure 3c), thus supporting the mixed phenology hypothesis (Figure 1, H4). Differences in the rates of phenological change along the latitudinal gradient were nonetheless significant only between Lepidoptera and Coleoptera (Wilcoxon W = 76233, *P*=0.025) (Table S3). Specifically, Lepidoptera had a mean difference in rate of VID variation between low (L_1_) and high (L_2_) latitudes of 0.49 d/y ± 0.01, significantly higher than the difference for Coleoptera (0.38 d/y ± 0.05). This finding suggests that Coleoptera may adapt faster than Lepidoptera to environmental changes along latitudinal gradients, consistent with estimates for Danish beetles and butterflies^38^.

Life-cycle voltinism was a key moderator of how insect phenology responded to climate change (Figure 3b, Table S4). Indeed, VID changed faster in univoltine (1 generation/yr) than multivoltine insects (>1 generation/yr; univoltine generation R_u_ = 0.29 ± 0.01, first generation R_m1_ = 0.19 ± 0.03; Wilcoxon W = 229365, *P*<0.001). This pattern is consistent with the findings of the phenological analysis of 130 lepidopteran species (including 39 multivoltine species) in the UK^3^, in that univoltine insects in our study advanced their phenologies less (-0.09 d/y ± 0.01) than did the first generation of multivoltine insects (-0.15 d/y ± 0.02). This finding suggests that multivoltine insects may adapt more quickly than univoltine insects to climate change, possibly because more generations have more chances to adjust their life history strategies^39^. Of the 223 species of multivoltine insects in our data set with sufficient records (>400 records for each generation), 70.4% had lower rates of shifts in VID for the first voltine pattern compared to the second (second generation R_m2_ = 0.38 ± 0.03, Wilcoxon V = 5977, *P*<0.001).

### Vegetation phenology is more sensitive than insect phenology to climate

The variation in VID could be attributed to the different directions and magnitudes of responses of vegetation and insect phenologies to the same environmental factors. We used statistical methods including partial correlation analysis, ridge regressions, and random forests (See Methods), to identify the likely environmental drivers of the phenological mismatch. The results of these analyses indicate that the average absolute sensitivity (|PC|, the partial correlation coefficient, and |RC|, the ridge regression coefficient) to environmental factors was generally higher for LUD than IOD, with six of the eight environmental factors differing highly significantly (*P*<0.001) (Figure 4a, b). This finding is consistent with long-term phenological studies in the northern Eurasian continent^18^, suggesting that the vegetation, as primary producers, was more adjustable than insects to environmental conditions. We obtained consistent outcomes for both PC and RC (Spearman ρ>0.73, *P*<0.001): VID tended to synchronize (PC_LUD-IOD_ and RC_LUD-IOD_ >0) with each unit increase in monthly evapotranspiration (ET), precipitation (PRE), wind speed (WIND), and soil-moisture content (SM). Conversely, VID shifted towards asynchrony (PC_LUD-IOD_ and RC_LUD-IOD_ < 0) (Figure 4c) for each unit increase in monthly soil humidity (HU), solar radiation (RAD), air temperature (TEM), and soil temperature (ST). The analysis of the trends of environmental factors over 34 years (Figure S2) found that the overall increases in RAD, TEM, and ST, and the decreases in ET and WIND likely led to the intensification of the asynchronous VID in Europe. The thermal indicators (TEM and ST) had the largest average impact on LUD (FI_ST_ = 0.16 ± 0.002, FI_TEM_ = 0.16 ± 0.002) and IOD (FI_ST_ = 0.13 ± 0.001, FI_TEM_ = 0.15 ± 0.001) (Figure 4c). These findings are in agreement with recent research elucidating the key influence of temperature on vegetation and insect phenologies^16,40^.

**Figure 4.**
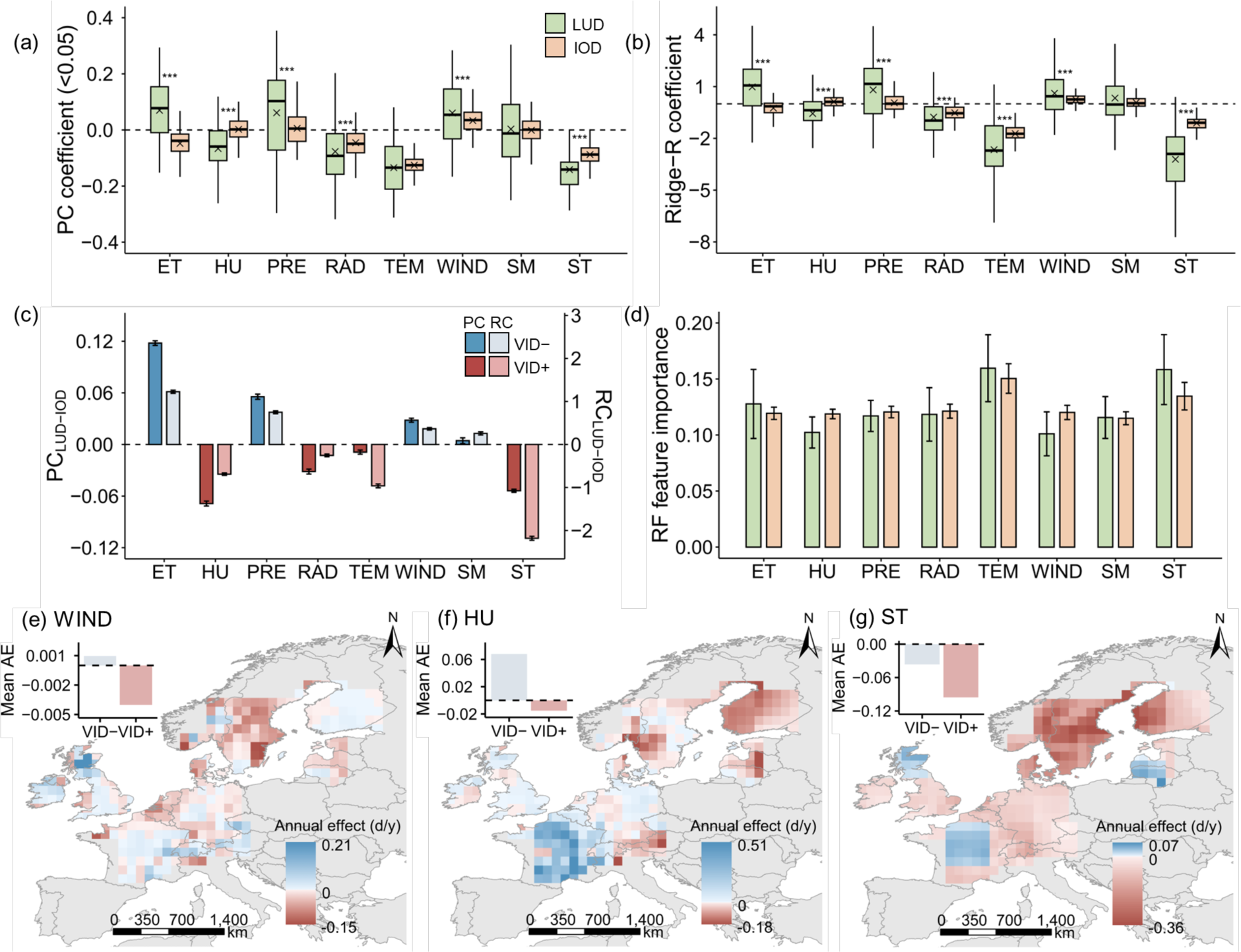
The asynchrony of the vegetation-insect phenological difference (VID) across Europe we report here has been primarily driven by the sensitivities of the leaf-unfolding date (LUD) and the insect occurrence date (IOD) to environmental factors. (a and b) LUD has a higher absolute sensitivity than IOD to environmental factors. (a) The partial correlation coefficient (PC) (only PC with *P*<0.05 were retained). (b) The ridge regression (Ridge-R) coefficients (RC); green refers to LUD, whereas orange refers to IOD. ***, *P*<0.001; **, 0.001≤*P*<0.01; *, 0.01≤*P*<0.05 by a Wilcoxon signed-rank test. (c) VID tends to synchronise with each unit increase in monthly evapotranspiration (ET), precipitation (PRE), average wind speed (WIND), and soil-moisture content (SM). VID shifts towards asynchrony for each unit increase in monthly relative humidity (HU), solar radiation (RAD), average temperature (TEM), and soil temperature (ST). Error bars represent ± S.E.; blue indicates VID tending towards synchrony (VID−: PC_LUD-IOD_ and RC_LUD-IOD_ >0), and red indicates VID tending towards asynchrony (VID+: PC_LUD-IOD_ and RC_LUD-IOD_ <0). (d) The thermal indicators (TEM and ST) are most important for LUD and IOD. Random forest (RF) feature importance, indicating the relative importance of each factor for LUD and IOD, with a collective sum of 1; error bars represent ± S.D. (e-g) WIND, HU, and ST are key drivers of the significantly greater phenological asynchrony in the VID+ than the VID− regions. The annual effect (AE) estimates the yearly impact of the environmental factors on VID, calculated as RC_LUD-IOD_ × R_ENV_ (annual rate of change of the environmental factors; Figure S4).

At the regional scale, the spatial differences of sensitivity of LUD and IOD to environmental factors, and trends of changes in environmental factors, collectively lead to an asynchronous intensification of VID along the latitudinal gradient. A total of 69% of VID values across Europe indicated a trend towards asynchrony (R_VID_ > 0; VID+), whereas only 31% showed a trend towards synchrony (R_VID_ < 0; VID−; Figure 2a). Further analyses identified a significant difference (*P*<0.05) in RC_LUD-IOD_ between the VID+ and VID− regions (Figure S3). An analysis of the annual effect of environmental factors found that the increases in ST and HU, and the decrease in WIND, led to a stronger phenological asynchrony in the VID+ regions compared to the VID− regions (Figures 4e-g; S4). The environmental sensitivities of LUD and IOD generally varied significantly along the latitudinal gradient (Figure S5), representing the differences in environmental adaptability between populations at high and low latitudes^41^.

## 3. Discussion

The extent and direction of changes in phenological synchrony are the outcome of intricate interactions between the life history strategies of plants and insects, their physiological characteristics, and external environmental stressors^30,37,42^. Here, we examined shifts in the phenology of 1,584 herbivorous insect species and the corresponding vegetation phenologies across Europe between 1982 and 2015, and found that the phenological synchrony between European vegetation and insects has gradually become more misaligned. This pattern was mostly attributed to more pronounced phenological advances for vegetation than insects, so maintaining the historical synchrony between insect and vegetation phenologies is likely to prove challenging going forward^37^. Notably, we found strong taxa- and life-cycle-specific effects on phenological shifts that are also in agreement with a study conducted in Ireland^43^. Finally, we found that the phenological mismatch across Europe was mostly due to increases in monthly radiation and air and soil temperatures, and decreases in evapotranspiration and wind speed.

One of the pivotal causes of the progressive misalignment between vegetation and insect phenologies was their different responses to external environmental conditions. Our study found that this asynchrony was not uniformly distributed but had distinct geographical patterns. Areas at higher latitudes have experienced a more pronounced decoupling of vegetation-insect phenological synchrony in recent decades. We found that this phenological asynchrony was not only associated with the pace and magnitude of climate change in these regions, but was also associated with the spatial variations in the sensitivities of vegetation and insect phenologies to different climatic factors, a finding in agreement with other reports in other regions across the world^15,44^. European vegetation has generally become more sensitive to environmental fluctuations compared to insects in the last 34 years. The phenological mismatch between vegetation and insects may continue to worsen over time, because climatic predictions project a future scenario of continued increases in global temperatures. Multispatial and temporal-scale analyses, such as ours, are imperative for understanding and forecasting this phenomenon due to the disparities across geographical regions and biological taxa^45^.

Open-access data sets containing data from the long-term monitoring of vegetation and insect phenologies have fortunately become more available. Such integrated data will aid in deepening our understanding of the interactive dynamics between vegetation and insects under climate change^46^. Approaching the analytical outcomes of these data sets with caution, however, is crucial due to inevitable taxonomic and biogeographical sampling biases^47^.

The phenological misalignment between vegetation and insects could have profound implications for food webs, ecosystem functions, and ecosystem services^9^. Vegetation-insect asynchrony may potentially strongly affect population structure, adaptability, and species niches^25,49,50^. We must identify the potential physiological and ecological mechanisms underlying the different phenological responses of vegetation and insects to enhance our ability to predict and mitigate the impacts of phenological asynchrony^45^. Future research should focus on the complex interactions within interspecific phenological networks and the potential impacts of both biotic and abiotic factors on phenological changes^24,51^. It should also pay close attention to high-latitude areas, where phenological changes are more pronounced^22^, as we found, which will aid in deepening our understanding of the unique patterns and consequences of phenological asynchrony in regions experiencing intensified climate change. These large-scale, integrative approaches will provide a robust scientific basis for ecological conservation, the management of resources, and the development of policies.

## Methods

### Satellite-derived data for spring vegetation phenology

The satellite-derived leaf unfolding date (LUD) was determined using the Normalized Difference Vegetation Index (NDVI). We thus accessed the GIMMS NDVI3g data set (http://poles.tpdc.ac.cn/en/data/), a product derived from the Advanced Very High-Resolution Radiometer (AVHRR)^52^. This data set has a temporal resolution of 15 days and a spatial resolution of 8 km^2^. We focused on pixels of vegetation in the temperate terrestrial ecosystems of Northern Hemisphere (>30°N) for 1982-2015, where the dynamics of vegetation phenology were strongly seasonal. We excluded pixels with an annual mean NDVI < 0.1 to reduce the effect of sparse vegetation^53^. We identified pixels with a continuous daily average temperature <0 °C for at least five days as the snow-contaminated phase and replaced them with the NDVI of the nearest snow-free date^54^ to minimise the impact of snow. We then smoothed the curves of daily NDVI time series using the Savitzky-Golay method^55^ to reduce the effect of noise from atmospheric disturbance. Finally, we extracted the average LUD using three methods to ensure the robustness of the results: (1) the dynamic-threshold method^56^, (2) the piecewise logistic function method^57^, and (3) the modified double-logistic function method^58^ (see details below).

The dynamic threshold method determines the threshold of LUD using the proportion of the smoothed NDVI annual time series as:

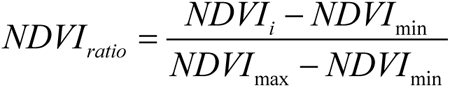

where *NDVI_i_* is the NDVI on the *i*-th day of a year (DOY), *NDVI*_max_ and *NDVI*_min_ are the annual peak and minimum NDVI values, respectively, and LUD is the date whenspring *NDVI_ratio_* first increases to 0.5.

The piecewise-logistic function method divides the smoothed curve of the NDVI time series into two periods, before and after the annual peak (*α*). This method then fits the daily curve using the piecewise logistic function:

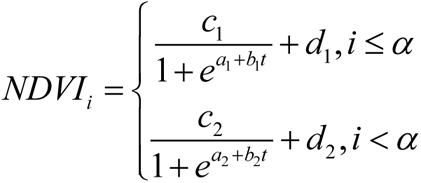

where *a* and *b* are fitting parameters, with *a* as the initial rate of increase in NDVI and *b* as the rate of change of NDVI with time, *c* is the amplitude of the change of NDVI change (i.e., the difference between the peak and background values), and *d* is the NDVI background value.

Finally, the modified double-logistic function method fits a smoothed curves of the NDVI time series using a double-logistic function:

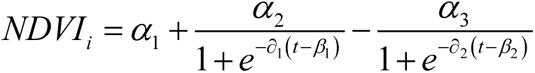

where *α*_1_ is the background NDVI, *α*_2_ is the difference between the summer peak NDVI and the spring minimum NDVI, and *α*_3_ is the difference between the summer peak NDVI and the autumnal minimum NDVI. ∂_1_ and ∂_2_ are the curvatures of the ascending and descending phases, respectively, and β_1_ and β_2_ are the average DOYs of vegetation greening and senescence, respectively.

### Records of occurrence of herbivorous insects across Europe

We used records of the occurrence of herbivorous insects from crowdsourced data sets to calculate insect phenology due to the limitations of precise spatial and temporal coverage and species representation of phenological records for herbivorous insects. We searched the Global Biodiversity Information Facility (GBIF, https://www.gbif.org/) for all occurrences of species of four insect orders (Hemiptera, Hymenoptera, Coleoptera, and Lepidoptera) across Europe (Lon -11° to 35°, Lat 34° to 71°) for 1982-2015. This search found a total of 2,937 insect species. We next excluded insects that primarily depended on predation, parasitism, or saprophagy during any stage of their life cycles. We also removed duplicate entries and records with inaccurate coordinates (uncertainty > 1 km^2^). The final database retained ≥8 million observations.

### Classification of insect phenological patterns

We classified the complex life histories of insects based on their degree of voltinism (i.e., the number of generations in one year). Variation in voltinism in insects can thus produce either unimodal or multimodal phenological patterns and is linked to important ecological processes such as population dynamics and community interactions^59^. The timing and peak period of occurrence of insect larvae and their adults may differ considerably^60^. These different developmental stages have distinct nutritional requirements and ecological effects, e.g. the herbivory of larvae and the pollinating behaviour of adults^61,62^. We removed records of occurrence with life-stage labels of egg, larval, or pupal stages (<1%) from our data set due to the lack of data to support modelling. We thus focused our modelling only on the mean date of occurrence of adults.

We calculated the probability density of species records for each DOY (Day of the year) to obtain the annual fluctuations of insect population and subsequently smoothed the probability-density curve using Gaussian filters^63^ and Savitzky-Golay filters to reduce noise. We next used the find_peaks function from the “scipy” Python package to determine whether a species had multimodal phenology (i.e., more than one population peak). We then refined the phenological patterns following a thorough comparison with both scientific and grey literature^6^. We found that 295 species had multimodal patterns of distribution (Figure S6). We also used the K-Means algorithm to classify the records of the occurrences of these species:

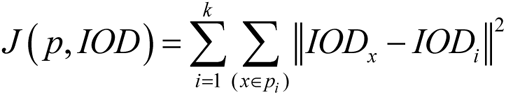

where *k* is the number of phenological patterns (number of clusters), *IOD_i_* is the mean date of the occurrence of insects for phenological pattern *p_i_*, and ‖*IOD_x_* − *IOD_i_*‖^2^ is the squared Euclidean distance between each insect occurrence date (*IOD_x_*) and the mean observed date (*IOD_i_*) of each group of phenological patterns. The K-Means algorithm was thus used to find the optimal phenological classification that minimises the mean squared error (MSE) of the objective function *J* _(_ *p*, *IOD*_)_ ^64^. We calculated k-means for *k* = 2 to the maximum number of peaks to determine the optimal number of patterns. We recorded the silhouette coefficient (SC)^65^ for each *k*. The number of clusters with the maximum SC was considered as the best number of patterns *k_opt_*. We removed outliers with residuals >3 S.D. during the clustering and removed phenological patterns with fewer than 400 records due to the uncertainty of the crowdsourced data in our data set. Finally, we identified 1,807 phenological patterns encompassing a total of 1,584 species (Figure S1).

### Spatiotemporal differences in the analysis of vegetation-insect phenology

We evaluated the overall trend of variation in vegetation phenology, insect phenology, and their differences (VID) over time and geographical locations. We applied a quantile-regression model to each of the 1,807 phenological patterns to ensure the robustness of our estimation results:

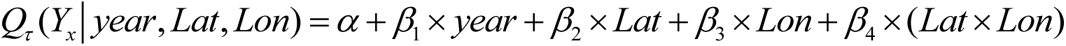

where, *Q_τ_* is the *_τ_* quantile, with *_τ_* = 0.5 set as the median. *Y_x_* is the response variable corresponding to observation record *x* (leaf unfolding date, *LUD_x_*; insects occurrence date, *IOD_x_*; or vegetation-insect difference, *VID_x_*). In this equation *α* is the intercept, and *β*_1_, *β*_2_, *β*_3_ and *β*_4_ are the coefficients for the year, latitude (Lat), longitude (Lon), and interaction term, respectively. We defined the rate of change of *Y_x_* relative to the predictor variables as the absolute value of *β_i_*; this rate of change was positive when the absolute value of the response variable increased between 1980 and 2015 and negative when it decreased.

We divided the phenological patterns into areas at high (>55°N) and low (≤55°N) latitudes for separate quantile-regression models. We calculated the trends of the four insect orders separately to avoid the dominance of Lepidoptera species (>70%) and to determine whether the changes in VID had significant geographical patterns, while also considering the limitations of spatial data. We also conducted spatial statistics to obtain local trends. We thus divided Europe into 5×5° geographical grids based on a grid analysis and used a moving-window algorithm^6,41^. Briefly, the algorithm moved by 1-4° in longitude or latitude, and the quantile-regression models of the phenological patterns for LUD, IOD, and VID were constructed grid-by-grid. We then averaged the results for the rate of change based on the 1×1° grid unit to minimise the variations in sampling public data across regions. We only retained the results from a single geographical grid that had at least 10 insect species, and each phenological pattern had a record of no less than 10 years^66^, to ensure the reliability and representativeness of the data.

## Analysis of the impact of environmental factors on vegetation and insect phenologies

### Data acquisition

We incorporated two comprehensive data sets to obtain long-term environmental data from 1982 to 2015 across Europe for elucidating the influence of environmental factors on the vegetation and insect phenologies (Table S5).

The E-OBS data set, a daily gridded data set of terrestrial observations for Europe, is based on the network of stations of the European Climate Assessment & Dataset (ECA&D) project^67^. This data set aggregates many types of data directly from the European National Meteorological and Hydrological Services. Its comprehensive nature makes it an indispensable data set for monitoring the climate across Europe^68^. We obtained meteorological data from the E-OBS 27.0e data set (accessible at https://www.ecad.eu/download/ensembles/download.php) on average air temperature (TEM), precipitation (PRE), average wind speed (WIND), average relative humidity (HU), and global solar radiation (RAD). These data, with a spatial resolution of 0.1×0.1°, were further processed to deduce monthly averages.

We incorporated soil parameters into our assessment in recognition of the pivotal interplay between soil and both vegetation growth and insect activity, most notably in the realms of nutrient cycling and habitat provisioning^69^, using the ECMWF Reanalysis v5 (ERA5)-Land reanalysis data set^70^. ERA5, unlike its predecessor, offers a consistent portrayal of the evolution of terrestrial variables over several decades, but at a heightened resolution. Specifically, we extracted data from the monthly ERA5-Land data set (accessible at https://cds.climate.copernicus.eu/#!/home) on soil temperature (ST) and soil-moisture content (SM). We also obtained evapotranspiration (ET) metrics from the data set, where negative values indicate evaporation and positive values indicate condensation, due to the integral role of transpiration in maintaining vegetation health and its consequential potential microclimatic impacts on insect populations^71^.

### Analytical strategy

We used a partial correlation analysis^72^ to separately identify the effects of the environmental factors on the vegetation and insect phenologies for understanding the underlying causes of changes in VID (TEM, PRE, WIND, HU, RAD, SM, ST, and ET in Figure 4). Environmental factors usually affect biological phenology in the months before its onset. Considering the variation in the times of insect observation, the environmental factors were detrended and standardized using linear regression to remove the multiyear trend^73^. The optimal preseason length was defined as the period during which phenology (IOD or LUD) and the environmental factors had the highest absolute correlation for each species, with the search extending up to six months before the average time of the phenology^4^. We then conducted a partial correlation analysis using the “pingouin” Python package, calculating the partial correlation coefficient (PC) and significance for each environmental factor with phenology, while controlling for other variables.

We used ridge regression (Ridge-R)^75^ and the random-forest method (RF)^76^ from the “sklearn” Python package to assess the contribution of the environmental factors to the shift in VID, minimising the influence of collinearity. Phenology (IOD or LUD) was the response variable, with environmental factors with optimal preseason times as the predictive variables. The regression coefficient (RC) in Ridge-R was used to assess environmental sensitivity, representing the number of days the phenology advanced or delayed per unit standardised change in the environmental factor. The feature importance in the RF method was evaluated by measuring the decrease in impurity (MSE) each time a feature was split^75^. We similarly used the moving-window algorithm to conduct the three methods at the regional scale to understand the spatial patterns of the impacts of the environmental factors on the variation of VID.

## Author Contributions

C.W. proposed the original idea. R.S-G. and W.H. designed the research framework. Y.H. processed all data, derived models, generated figures, and drafted the manuscript. R.S.-G. guided its execution. All authors contributed to the writing of this study.

## Acknowledgments

C.W., W.H., and Y.D. were supported by National Key R&D Program of China (2023YFB3906200) and National Natural Science Foundation of China (42125101, 42071320). Y.H. was supported by the China Scholarship Council (202204910383). R.S.-G. was supported by NERC Pushing the Frontiers (NE/X013766/1).

