## Supplementary Materials for "Climate change has desynchronized insect and vegetation phenologies across Europe"

**Table of contents:**

| | |
|---|---|
| **Appendix S1: Phenological data and environmental data** | **P. 2** |
| Figure S1. Phenological data | P. 2 |
| Table S5. Environmental data | P. 3 |
| Original phenological and environmental data | (Additional files) |
| **Appendix S2: Extended results** | **P. 5** |
| Table S1. LUD, IOD, and VID shift rate | (Additional form) |
| Table S2. LUD, IOD, and VID at different spatial scales | P. 5 |
| Table S3. VID across four insect orders | P. 6 |
| Table S4. VID and IOD as a function of voltinism | P. 9 |
| Figure S2. Temporal trends in environmental drivers | P. 11 |
| Figure S3. LUD-IOD as a function of environmental drivers | P. 13 |
| Figure S4. Annual effect of environmental drivers | P. 14 |
| Figure S5. Spatial patterns of environmental sensitivity | P. 16 |
| Figure S6. IOD phenological patterns (and additional files) | P. 17 |
| **References** | **P. 18** |

### Appendix S1: Phenological data and environmental data

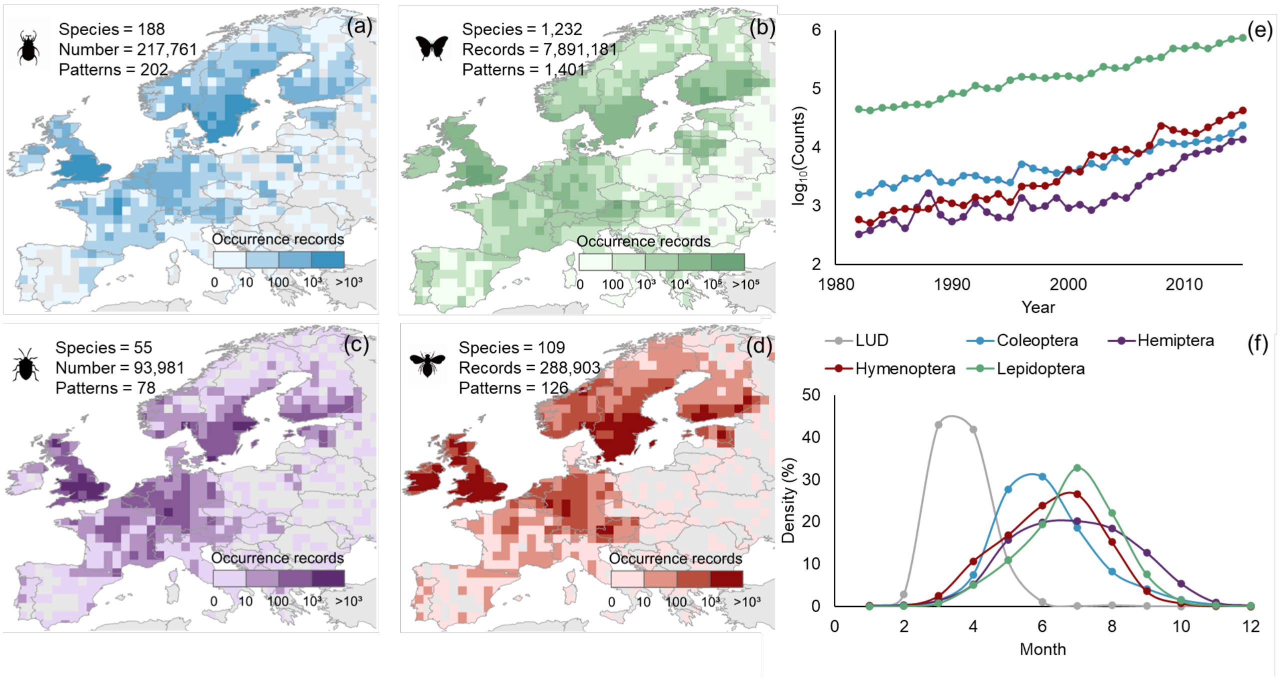

**Figure S1.** Phenological data of vegetation and insects in Europe from 1982 to 2015. (a-d) Occurrence records of four insect orders from GBIF (https://www.gbif.org/) after data cleaning; (a) Coleoptera (b) Lepidoptera (c) Hemiptera, and (d) Hymenoptera; The colour gradient from light to dark represents an increase in occurrence. (e) Annual count data of the occurrence of the four insect orders, showing an increasing trend over the last decades. (f) The monthly occurrence density statistics for both leaf-unfolding date (LUD) of vegetation and the four insect orders, illustrating that the average LUD of vegetation typically occurs earlier than the peak activity of the examined herbivorous insects.

**Table S5.** Environmental data of E-OBS[1] and ERA5-LAND[2] were used in this study to examine the drivers of phenological mismatch between insects and vegetation across Europe between 1982 and 2015. The daily data for E-OBS is aggregated into monthly average data. Within a given geographic grid, there must be at least ≥10 days of non-missing values per month; missing values are ignored in the calculations.

| Data source | Spatial resolution | Temporal resolution | Variable name in original data source | Acronym used in our study | Unit | Description |
|---|---|---|---|---|---|---|
| **E-OBS 27.0e** | 0.1° | Daily | TG (Mean temperature) | TEM | ℃ | Mean temperature recorded near the ground level, typically at a height of 2 m. |
| | | | RR (Precipitation sum) | PRE | mm | Total precipitation, including rain, snow, and hail, quantified by the depth of water it would form per square metre ($m^2$). |
| | | | FG (Mean wind speed) | WIND | m/s | Mean wind speed measured at an elevation of 10 m. |
| | | | HU (Mean relative | HU | % | Mean humidity close to the |

| | | | | | | |
|---|---|---|---|---|---|---|
| | | | humidity) | | | ground, usually measured at around 2 m above the surface. |
| | | | QQ (Global radiation) | RAD | $W/m^2$ | Mean flux of shortwave radiation measured at the Earth's surface. |
| **ERA5-LAND** | 0.1° | Monthly | soil_temperature_level_1 | ST | K | Soil temperature in the top soil layer (0 to 7 cm). |
| | | | volumetric_soil_water_layer_1 | SM | Volume fraction | Moisture content in the top soil layer (0 to 7 cm). |
| | | | total_evaporation_sum | ET | m of water equivalent | Total water evaporated from the Earth's surface over time, with negative values indicating evaporation and positive values signifying condensation. |

### Appendix S2: Extended results

**Table S2.** Leaf-unfolding date of vegetation (LUD), insect occurrence date (IOD), and vegetation-insect phenological difference (VID) annual shift rates at the regional scale ($L_1$≤55°N or $L_2$>55°N) and continental scale (Overall: $L_1$ and $L_2$). The Wilcoxon W value represents the outcome of the Wilcoxon rank-sum test, a non-parametric unpaired mean test. The Wilcoxon V value corresponds to the results from the Wilcoxon signed-rank test, a non-parametric paired mean test. Bolded *P*-values are significant at <0.05.

| | **VID change rate (Mean ± S.E. d/yr)** | **LUD change rate (Mean ± S.E. d/yr)** | **IOD change rate (Mean ± S.E. d/yr)** | **Wilcoxon V (LUD and IOD)** |
|---|---|---|---|---|
| **$L_1$ (≤55°N)** | -0.02 ± 0.004 | -0.19 ± 0.005 | -0.16 ± 0.004 | 4246 (***P*=0.001**) |
| **$L_2$ (>55°N)** | 0.38 ± 0.007 | -0.42 ± 0.005 | 0.02 ± 0.003 | 0 (***P*<0.001**) |
| **Overall ($L_1$ and $L_2$)** | 0.28 ± 0.009 | -0.37 ± 0.006 | -0.09 ± 0.007 | 199622 (***P*<0.001**) |
| **Wilcoxon W ($L_1$ and $L_2$)** | 1191 (***P*<0.001**) | 19126 (***P*<0.001**) | 2302 (***P*<0.001**) | - |

**Table S3.** Vegetation-insect phenological difference (VID) annual shift rates of four insect orders at the regional scale ($L_1$≤55°N or $L_2$>55°N) and continental scale (Overall: $L_1$ and $L_2$). The Wilcoxon W value represents the outcome of the Wilcoxon rank-sum test, a non-parametric unpaired mean test. The Wilcoxon V value corresponds to the results from the Wilcoxon signed-rank test, a non-parametric paired mean test. The abbreviations for Lepidoptera, Hemiptera, Hymenoptera, and Coleoptera are *Le*, *He*, *Hy*, and *Co*, respectively. Bolded *P*-values are significant at <0.05.

| | **Lepidoptera (Mean ± S.E. d/yr)** | **Hemiptera (Mean ± S.E. d/yr)** | **Hymenoptera (Mean ± S.E. d/yr)** | **Coleoptera (Mean ± S.E. d/yr)** |
|---|---|---|---|---|
| **$L_1$ (≤55°N)** | 0.01 ± 0.01 | 0.02 ± 0.07 | -0.26 ± 0.06 | -0.12 ± 0.04 |
| **Wilcoxon W (Orders at $L_1$)** | *Le & He:* 36379 (*P*=0.966)<br>*Le & Hy:* 31143 (***P*<0.001**)<br>*Le & Co:* 69103 (***P*<0.001**) | *He & Hy:* 3608 (***P*=0.003**)<br>*He & Co:* 4272 (***P*=0.045**) | *Hy & Co:* 7662 (***P*=0.046**) | - |
| **$L_2$ (>55°N)** | 0.50 ± 0.01 | 0.58 ± 0.07 | 0.20 ± 0.06 | 0.26 ± 0.04 |

| | | | | |
|---|---|---|---|---|
| **Wilcoxon W (Orders at $L_2$)** | *Le & He:* 38695 (*P*=0.361) *Le & Hy:* 30095 **(*P*<0.001)** *Le & Co:* 54749 **(*P*<0.001)** | *He & Hy:* 3827 **(*P*<0.001)** *He & Co:* 3277 **(*P*<0.001)** | *Hy & Co:* 6754 (*P*=0.811) | - |
| **Wilcoxon V ($L_1$ and $L_2$)** | 30798 **(*P*<0.001)** | 290 **(*P*<0.001)** | 647 **(*P*<0.001)** | 2351 **(*P*<0.001)** |
| **Overall ($L_1$ and $L_2$)** | 0.34 ±0.01 | 0.16 ± 0.05 | 0.15 ± 0.04 | 0.04 ± 0.03 |
| **Wilcoxon W (Overall trend of orders)** | *Le & He:* 71045 **(*P*<0.001)** *Le & Hy:* 110916 **(*P*<0.001)** *Le & Co:* 213020 **(*P*<0.001)** | *He & Hy:* 4752 (*P*=0.693) *He & Co:* 9230 **(*P*=0.026)** | *Hy & Co:* 10005 **(*P*=0.001)** | |

| | | | | |
|---|---|---|---|---|
| **Gradient differences ($L_2$ - $L_1$)** | 0.49 ± 0.01 | 0.56 ± 0.09 | 0.46 ± 0.08 | 0.38 ± 0.05 |
| **Wilcoxon W (Gradient differences among orders)** | *Le & He:* 39089 ($P$=0.288) *Le & Hy:* 46263 ($P$=0.884) *Le & Co:* 76233 (**$P$=0.025**) | *He & Hy:* 3009 ($P$=0.445) *He & Co:* *4335* ($P$=0.063) | *Hy & Co:* 6128 ($P$=0.332) | - |

**Table S4.** Vegetation-insect phenological difference (VID) and insect occurrence date (IOD) annual shift rates of three voltinism levels at continental scale (Overall: $L_1$ and $L_2$). The Wilcoxon W value represents the outcome of the Wilcoxon rank-sum test, which is a non-parametric unpaired mean test. The Wilcoxon V value corresponds to the results from the Wilcoxon signed-rank test, indicative of a non-parametric paired mean test. The abbreviations for univoltine, multivoltine first generation, multivoltine second generation are U, $M_1$, $M_2$, respectively. Bolded *P*-values are significant at <0.05.

| | **Univoltine (Mean ± S.E. d/yr)** | **Multivoltine first generation (Mean ± S.E. d/yr)** | **Multivoltine second generation (Mean ± S.E. d/yr)** |
|---|---|---|---|
| **VID ($L_1$ and $L_2$)** | 0.29 ± 0.01 | 0.19 ± 0.03 | 0.38 ± 0.03 |
| **Wilcoxon W (U & $M_1$/ $M_2$) Wilcoxon V ($M_1$ & $M_2$)** | U & $M_1$: 229365 (***P*<0.001**) U& $M_2$: 122802 (***P*<0.001**) | $M_1$ & $M_2$: 5977 (***P*<0.001**) | - |
| **IOD ($L_1$ and $L_2$)** | -0.09 ± 0.01 | -0.15 ± 0.02 | 0.04 ± 0.02 |
| **Wilcoxon W (U & $M_1$/ $M_2$) Wilcoxon V** | U & $M_1$: 225974 (***P*<0.001**) | $M_1$ & $M_2$: 3859 (***P*<0.001**) | - |

| | |
|---|---|
| **($M_1$ & $M_2$)** | U& $M_2$:<br>94511<br>(***P*<0.001**) |

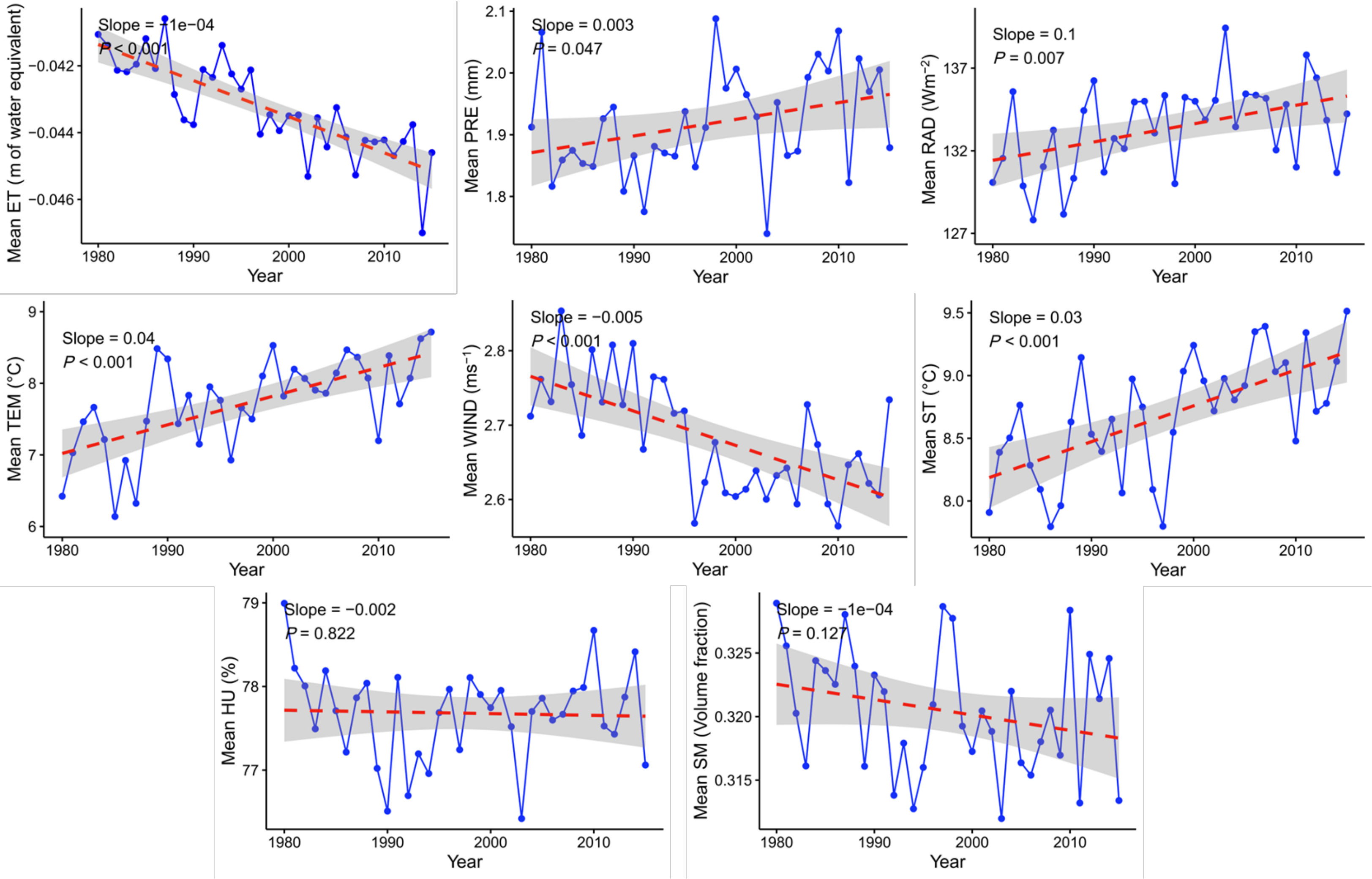

**Figure S2.** Trends in environmental factors over the 34 years of our study. A set of linear regressions were performed using annual mean data from Europe (Lon: -11° to 35°, Lat: 34° to 71°), where blue dots and lines represent the annual averages; the red dashed line represents the linear regression trend line, and the shaded grey area indicates the standard deviation of the trend line. Environmental factors include: ET: evapotranspiration, HU: average relative humidity, PRE: precipitation, RAD: solar radiation, TEM: average temperature, WIND: average wind speed, SM: soil-moisture content, ST: soil temperature (See Table S5).

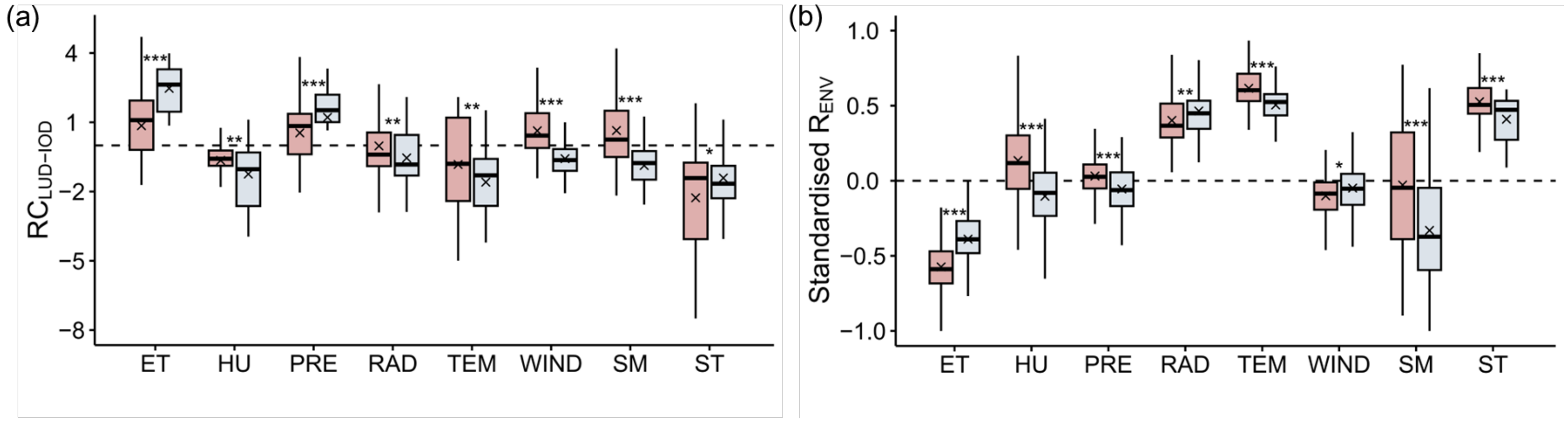

**Figure S2.** The differences in the environmental sensitivity of vegetation and insects, and environmental changes between synchronisation (VID−) and asynchronisation (VID+) regions. (a) The difference in environmental sensitivity between leaf-unfolding date of vegetation (LUD) and insect occurrence date (IOD) ($RC_{LUD\text{-}IOD}$) in the VID+ (red) and VID− regions (blue). (b) Standardised rate of environmental change during the examined 1982-2015 period. To account for the differences in units between different environmental factors, the data were standardised using the method of maximum absolute value normalisation, which preserves the original data's positive and negative values. ***: $P<0.001$; **: $0.001\leq P<0.01$; *: $0.01\leq P<0.05$ as per Wilcoxon signed-rank tests. The environmental drivers are: ET: evapotranspiration, HU: average relative humidity, PRE: precipitation, RAD: solar radiation, TEM: average temperature, WIND: average wind speed, SM: soil-moisture content, ST: soil temperature (See Table S5).

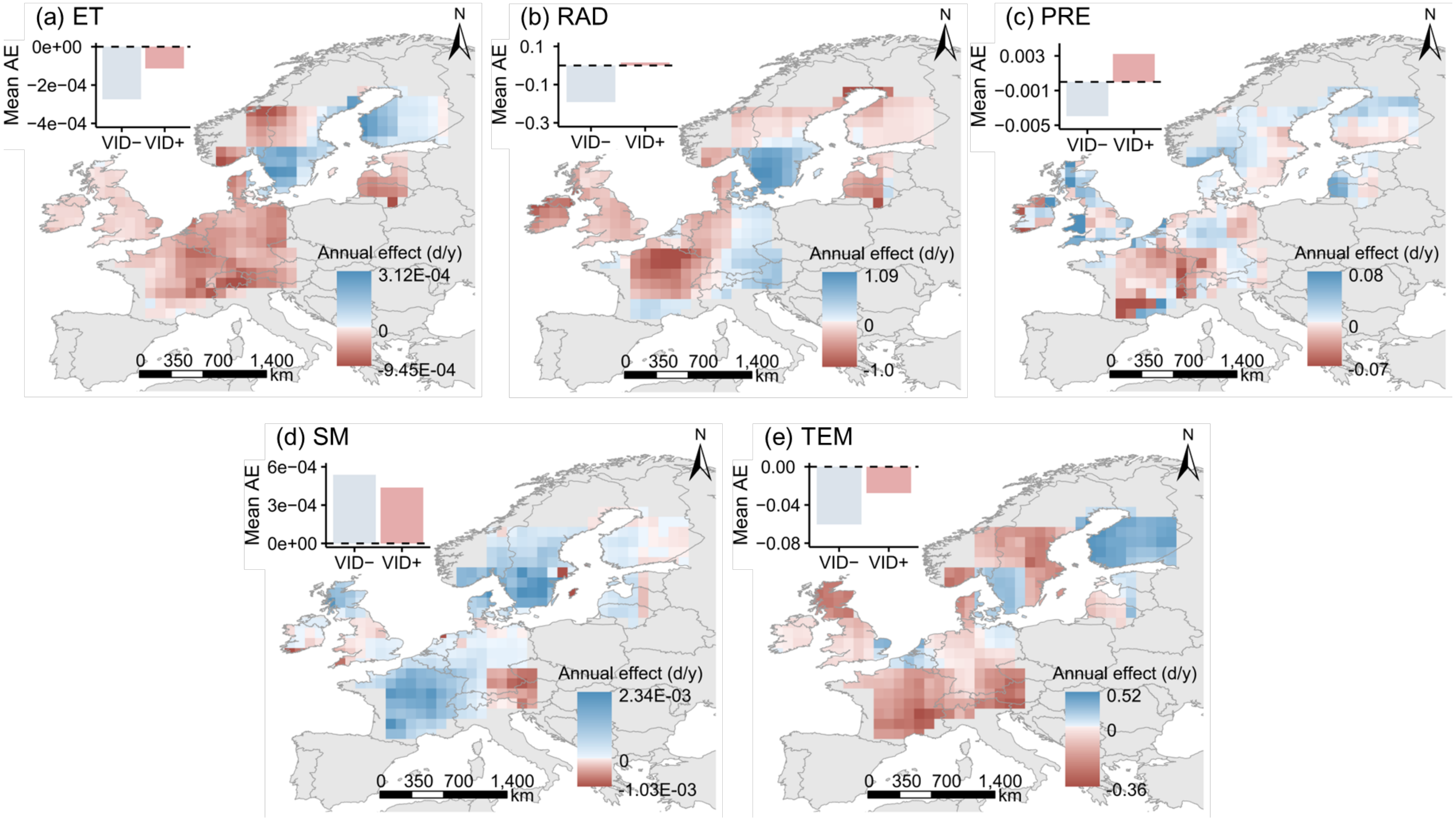

**Figure S4.** Annual effect (AE) estimates the yearly impact of environmental factors on the vegetation-insect phenological difference (VID). AE is calculated as $RC_{LUD\text{-}IOD} \times R_{ENV}$. Here, $RC_{LUD\text{-}IOD}$ represents the differences in environmental sensitivity of vegetation and insects, calculated using the ridge regression method. For more details on this calculation, refer to the 'Analytical strategy' section in the main article. $R_{ENV}$, on the other hand, denotes the annual change rate of the environmental factors, calculated as the linear regression coefficients of their annual average values at each grid point. Environmental factors include: ET: evapotranspiration, PRE: precipitation, RAD: solar radiation, TEM: average temperature, SM: soil-moisture content (See Table S5).

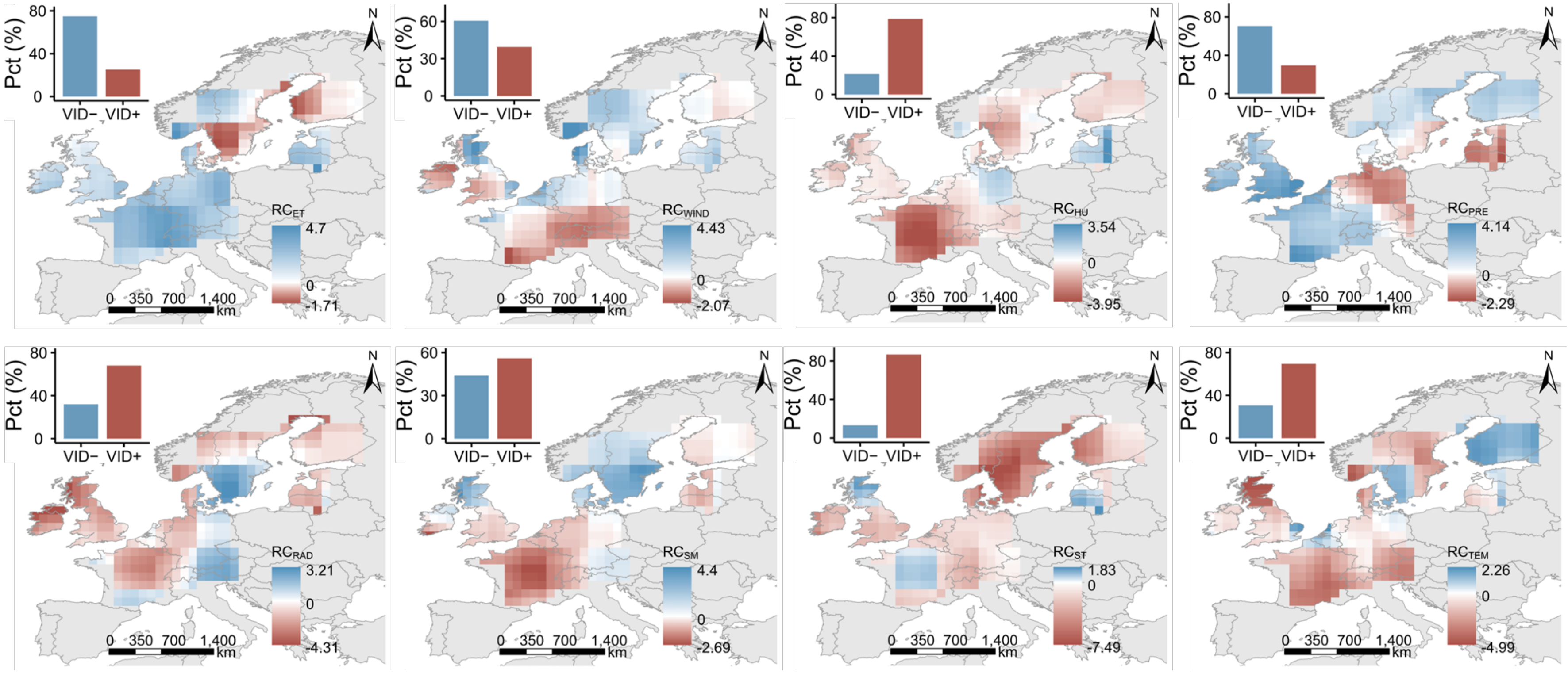

**Figure S5.** Spatial patterns of vegetation and insect phenology sensitivity differences ($R_{LUD-IOD}$) to environmental factors; The darker the colour (either red or blue), the higher the degree of asynchrony (Red; VID+) and synchrony (Blue; VID−); the inserted bar chart at the top-right of each panel represents the proportion of grid cells tending towards asynchrony and synchrony. Environmental factors include: ET: evapotranspiration, HU: average relative humidity, PRE: precipitation, RAD: solar radiation, TEM: average temperature, WIND: average wind speed, SM: soil-moisture content, ST: soil temperature (See Table S5).

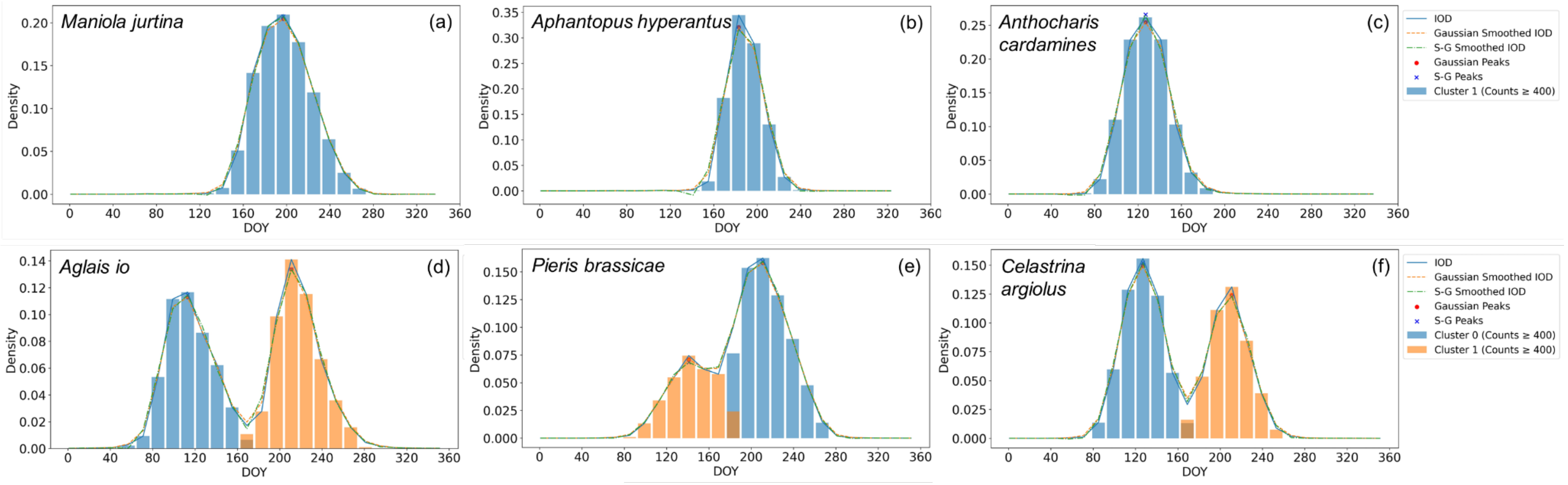

**Figure S6.** Classification of insect phenological patterns. (a-c) single phenological pattern, i.e., one generation per year, exemplified by *Maniola jurtina*, and *Aphantopus hyperantus*, *Anthocharis cardamines*. Peaks identified from the occurrence record density curve, after Savitzky-Golay and Gaussian filtering, are: 1. (d-f) single phenological insects, i.e., two or more generations per year, exemplified by *Aglais io*, *Pieris brassicae*, and *Celastrina argiolus.* Peaks identified from the occurrence record density curve, after Savitzky-Golay and Gaussian filtering, are: ＞1. Only phenological patterns with more than 400 occurrence records are retained. Further details are presented in the Methods section. Figures depicting the phenological patterns of other species are available in the Additional Files.
